# Ground-nesting bee survival after soil translocation and disturbance

**DOI:** 10.64898/2026.10.04.756489

**Authors:** Anne-Christine Mupepele, Alexandra-Maria Klein, Felix Fornoff

## Abstract

Ground-nesting wild bees require suitable soil conditions for their below-ground development and overwintering life stages. Understanding how soil translocation and disturbance affect individuals and the following generations of bees is important for identifying conservation measures for ground-nesting wild bees. We investigated the effects of soil translocation and disturbance on the ground-nesting solitary spring mining bee *Colletes cunicularius*. In a replicated controlled design, we compared the effects of soil translocation with different degrees of disturbance on the number of surviving bees and the emergence time. We found that soil translocation and disturbance reduced bee survival but did not result in complete mortality, with some individuals surviving even under severe soil disturbance. There was no evidence that soil disturbance affected the timing of spring emergence, suggesting that disturbance is unlikely to alter the timing of nest construction and, consequently, the production of the next generation. Our results indicate that translocating soil containing overwintering bees could be a promising conservation measure to compensate for habitat loss caused by land conversion.

## Introduction

The conservation of wild bees depends on the availability of suitable habitat conditions, including resources, such as nutritional flowers and nesting sites (Ellis *et al*., 2026). For nutrition, they depend on floral resources with adult bees primarily feeding on nectar, whereas larvae develop on a pollen-nectar mixture. In addition to floral resources, wild bees require nesting sites. Depending on the species, nests are constructed in pre-existing cavities aboveground, or in the soil belowground. More than half of all wild bee species are belowground-nesting and establish their nests in the soil (Michener, 2007; Harmon-Threatt, 2020). Adult female bees excavate nests in the soil and provision brood cells with pollen before laying an egg in each cell. The adult aboveground life stage is typically short-lived, and many univoltine solitary bee species have flight periods lasting only a few weeks each year (Antoine and Forrest, 2021; Michener, 2007; Danforth *et al*., 2019). For the remainder of their life cycle, bees persist belowground as developing offspring or dormant individuals within their nests. Consequently, belowground conditions play a crucial role in the conservation of wild bee populations.

Human activities alter landscapes and modify both aboveground vegetation and belowground soil conditions through land-use practices (Guo and Gifford, 2002; Kowalchuk *et al*., 2002; Borrelli *et al*., 2017; Haddaway *et al*., 2017). Land-use impacts on bees have been studied with a focus on alternating floral resources or aboveground-nest availability, with less attention given to belowground-nesting bees and their soil requirements (Orr *et al*., 2022; Hellerich *et al*., 2025; Tschanz *et al*., 2025; Thapa-Magar *et al*., 2022). Human-induced soil disturbance occurs through agricultural practices such as ploughing and through urban land-use pressures that increase soil sealing for buildings. Addressing this requires targeted translocation and restoration to maintain habitat connectivity that sustains bee populations across landscapes (European Commission, 2023; Ralls *et al*., 2018; Rayfield *et al*., 2011; Tischendorf and Fahrig, 2000). Ploughing is used to prepare fields for new crops and potentially influences bees, e.g. by exposing over-wintering individuals to harsh conditions (Ullmann *et al*., 2020; Hellerich *et al*., 2026; Roger-Estrade *et al*., 2010; de Graaff *et al*., 2019; Yin *et al*., 2022; Gruss *et al*., 2023; Boetzl *et al*., 2022). Bees nest mostly in the higher soil layers, and since ploughing commonly affects the top 20 to 30 cm, nesting bees are likely impacted (Harmon-Threatt, 2020; Roulston and Goodell, 2011; Williams *et al*., 2010; Ullmann *et al*., 2016; Tschanz *et al*., 2022).

Besides ploughing in agricultural landscapes, soils can also be sealed, for example through the construction of roads and buildings in urban areas. Soil sealing leads to a complete destruction of the habitat for belowground-nesting bees (Geisen *et al*., 2019). In such cases a translocation of populations and communities may be considered (Pérez *et al*., 2012; International Panel on Biodiversity and Ecosystem Services (IPBES), 2016; Lecocq *et al*., 2017). Translocation can be used to conserve populations threatened by habitat destruction, e.g. from construction, or to establish new populations in areas where they have previously been lost, thereby restoring connectivity across otherwise simplified and homogenized landscapes (Parker and Potter, 1974; Osgood, 1974; Donovan *et al*., 2010; Ahlbäck Widenfalk *et al*., 2018). Bee translocations can be conducted either during the adult aboveground stage or during immature developmental stages, when individuals are not yet flying. Adult translocations involve actively moving individuals to new locations, but success has often been limited because many bee species are philopatric, tending to establish nest sites at the same location where they themselves hatched and showing reluctance to relocate once nest construction has begun (Osgood, 1974; Parker and Potter, 1974; Cane, 1997; Kierat and Filipiak, 2017). Alternatively, bees can be translocated during their immature stages. This approach is relatively straightforward for aboveground-nesting species, but more challenging for belowground-nesting bees. For belowground-nesting bees, it requires either excavating nests or relocating intact soil blocks containing brood cells (Cronin, 2001; Gathmann and Tscharntke, 2002; Noll, 1931; Parker and Potter, 1974; Donovan *et al*., 2010). A potential advantage of soil-block translocation is that it can simultaneously move other soil-dwelling invertebrates and plant seeds, thereby transferring more components of the local ecosystem rather than only the target bee species (Vécrin and Muller, 2003; Bruelheide, 2003; Donovan *et al*., 2010; Smith *et al*., 2012). Successful translocations depend on multiple factors and translocation methods should be evaluated before implementation.

Soil disturbance, whether caused by soil turnover or translocation, plays an important role as restoration goals increasingly compete with land-use demands driven by a growing human population (Mupepele *et al*., 2021). Soil disturbance can affect belowground-nesting bees (hereafter ground-nesting), reducing the individual survival probability or the emergence time of adults (Hayes and LÓpez-Martínez, 2021; Pitts-Singer and James, 2005). In this study, we investigate the effects of soil disturbance on the solitary ground-nesting spring mining bee *Colletes cunicularius*. This species forms large nesting aggregations that allow reliable estimation of survival rates. *C. cunicularius* is univoltine, making populations particularly vulnerable to annual disturbances such as ploughing, as population losses cannot be compensated by a second generation within the same year. We investigate bee survival among different soil disturbance treatments and assess whether survival differs between males and females. Furthermore, we examine whether soil disturbance affects emergence time hypothesizing that bees originating from disturbed soils emerge earlier than those from undisturbed soils. Understanding both survival and phenological responses is essential for evaluating the effect of soil disturbance and to assess the effectiveness of soil translocation as a potential conservation measure, restoring connectivity through targeted reintroductions of populations.

## Methods

### Study site and experimental design

The study was conducted in Freiburg im Breisgau, a city in the southwest of Germany. A large nest aggregation of the solitary ground-nesting *C. cunicularius* has been observed for several years at the city’s exhibition site where the parking area is bordered by drainage ditches with sandy slopes (first observations by the authors in 2019. Geographic Coordinates (WGS84 from Google Maps): 48.01869481 N; 7.826365652 E, Fig. 1).

**Figure 1.**
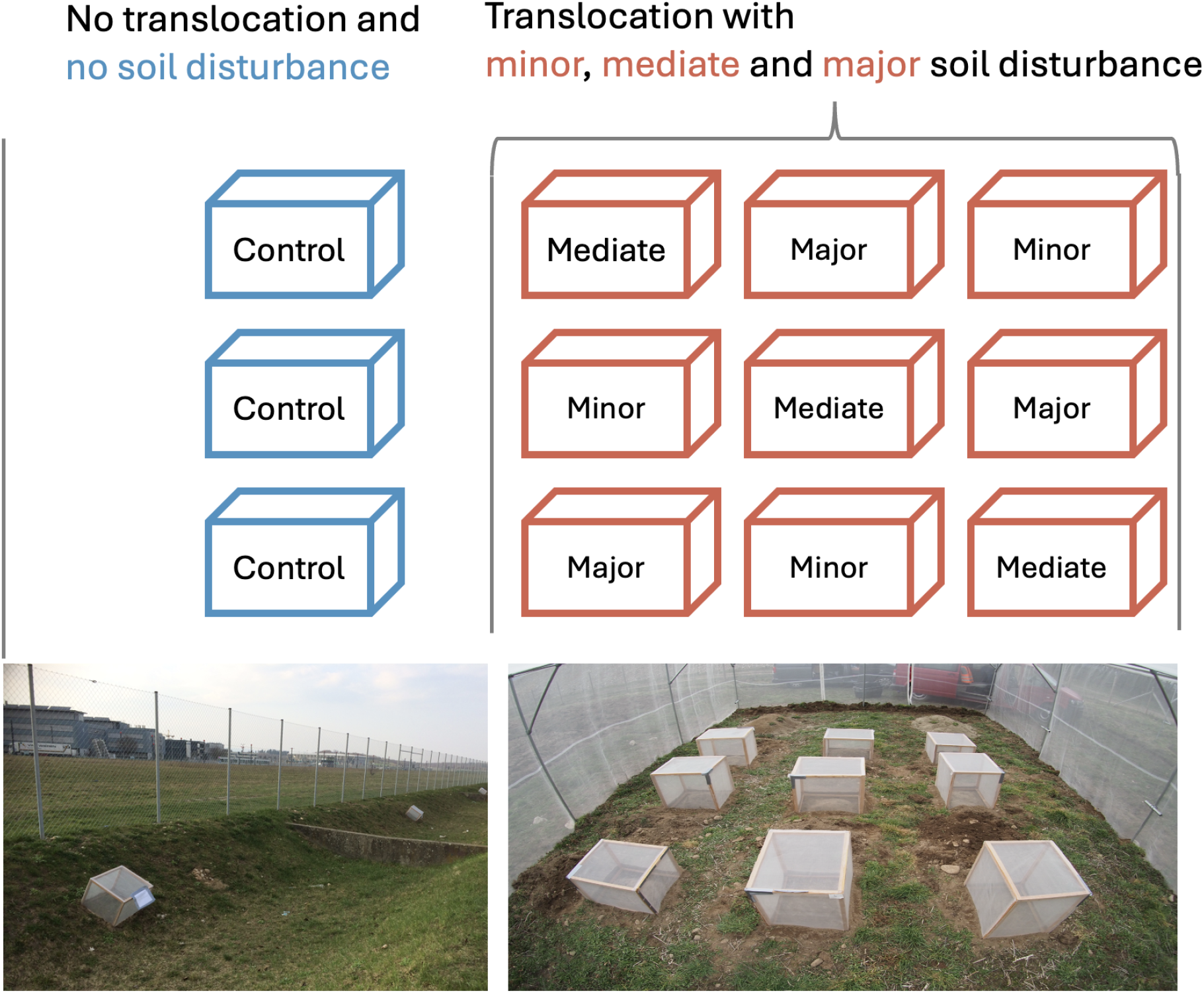
The study design consists of a control site, i.e. three emergence traps placed over a *C. cunicularius* nest aggregation without soil disturbance (left picture). For the three different translocation treatments, nine soil cubes were excavated at the nest aggregation and translocated to a nearby experimental site of the University of Freiburg. Translocated soil cubes were subject to three different soil disturbance treatments (minor to major). Emergence traps were also placed over each translocated soil cube (bottom right picture).

To test soil disturbance impacts on the survival and emergence time of the bees, we excavated nine 40 × 40 × 40 cm (64 000 cm^3^) soil cubes (Fig.1, Fig. S1) from the drainage ditch at the exhibition site. We excavated soil cubes on the 3rd of March 2021 before bees would end hibernation and start emerging from the soil. The disturbed soil samples were translocated in boxes by car to the experimental sites of the University of Freiburg at approximately one kilometer of the exhibition site (Fig. S1).

The disturbed and translocated soil was used for three different disturbance treatments: First, a ‘minor disturbance’ that involved that soil cubes were gently transferred to the experimental site and buried again to the same depth without turning the soil over. Second, a ‘mediate disturbance’ following the same procedure than the first treatment, but this time including a turnover of the soil. Soil was once turned upside down by pouring the 40 x 40 x 40 cm soil cubes from one meter height to the ground (Fig. S1). Third, a ‘major disturbance’ was subject to soil turnover by three times pouring the 40 x 40 x 40 cm soil cube from 1 m height to the ground and mimicking heavy machinery with four spade pushes. Undisturbed plots at the exhibition site served as control plots and were not subject to translocation or any other soil disturbance (Fig. 1).

Each treatment was replicated three times, and emergence traps were installed over the translocated and disturbed soil cubes, as well as at the exhibition site to collect control samples (Fig.1). Emergence traps were constructed from a wooden frame and covered with insect mesh on all sides except the bottom, enclosing an area of 66 x 56 cm (Fig. 1). Traps were checked on a daily basis and bees were collected alive from the net of the emergence traps where they would usually sit after emerging. All bees were sex-determined and identified to species levels.

### Statistical Analysis

#### Bee survival response to soil disturbance (H1)

Bee survival was assessed by counting the number of bees per emergence trap. Since emergence traps covered an area of 66 x 56 cm, whereas translocated soil cubes measured only 40 × 40 cm, bee counts of the control site were standardized to the same surface area than the treatment cubes represented. After standardization, bee counts were used as response variable in the linear models.

To test the effect of soil-disturbance treatments on bee survival, we applied an Analysis of Variance (ANOVA). We additionally included sex (female versus male bees) and the interaction between sex and treatment as predictor variables to assess whether treatment effects differed between sexes. Model assumptions, such as normality and homogeneity of variance, were checked a priori (Appendix S6). Likelihood ratio tests were used for model selections, starting with the full two-factorial ANOVA considering treatment, sex and the interaction as predictor and the standardized number of bees per trap as response variable. Subsequent posthoc comparisons were used in case of detecting an influence in the ANOVA (Tukey Honest Significant Differences, (Miller, 1981; Yandell, 1997)).

#### Sensitivity Analysis

Our disturbance treatments consisted of soil cubes excavated to a depth of 40 cm, with most brood cells found at depths of 20 to 30 cm (personal observation). At control sites, where soil remained undisturbed, bees could have potentially nested at greater depths. Given that *C. cunicularius* nests at a maximum depth of 60 cm (Harmon-Threatt, 2020), we conducted a sensitivity analysis in which control samples were standardized to a depth of 60 cm. This meant that control site samples were standardized not only to the reduced surface area of the treatments but also to their sampling depth. Accordingly, control site samples were rescaled from a volume of 66 × 56 × 60 cm to match the 40 × 40 × 40 cm treatment volume.

#### Emergence time of bees in response to soil disturbance (H2)

Emergence date was analysed on a Julian day scale, using generalized linear mixed models to test the effects of treatment and sex, with trap included as a random effect and a Gamma error distribution with a log link (‘glmmTMB’ package (Brooks *et al*., 2017; McGillycuddy *et al*., 2025)). The best model was selected using residual diagnostics of the ‘DHARMa’ package (Hartig, 2024) and likelihood-ratio-tests of the ‘stats’ package (R Core Team, 2025).

## Results

We sampled 212 bee individuals of the spring mining bee, which emerged from the soil between 6th of March and 5th of April 2021. Males and females were approximately equally abundant (115:97). There were no other bee species emerging from the soil, besides one female of *Sphecodes albilabris*, the cuckoo bee species of *C. cunicularius* that was emerging from one of the soil cubes in the minor disturbance treatment and was not included in the analyses.

### Soil disturbance influences bee survival

Soil disturbance had an impact on the number of surviving bees. Fewer bees emerged and thus survived if a disturbance took place in comparison to the undisturbed control sites (ANOVA: F=12, df=3, p*<*0.001, Fig 2). There was no further evidence for differences between the disturbance treatments (minor, mediate and major disturbance, Table S2, Appendix S6) nor for differences between male and female bees surviving as a consequence of soil disturbance (Table S1, Appendix S6).

**Figure 2.**
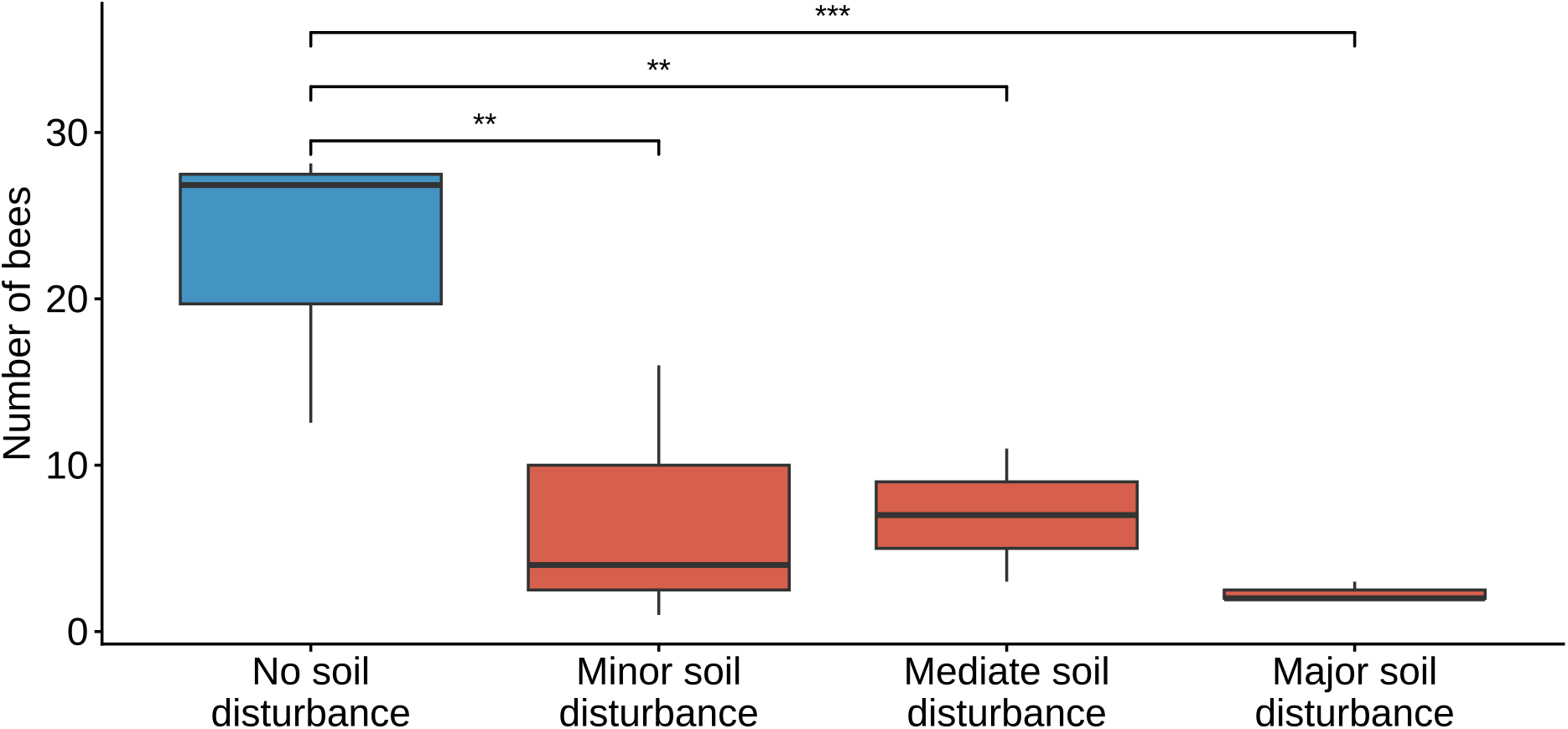
: Number of bees (standardized bee counts) in the four soil disturbance treatments: No soil disturbance (control), minor, mediate and major soil disturbance.

### Sensitivity analysis

The results were robust and the negative effect of soil disturbance on bee survival remained, even under the conservative assumptions that bees may have nested up to a depth of 60 cm (ANOVA F-statistic=6.12, df=3, p= 0.0042, Appendix S6).

### No evidence for an influence on emergence time

There is no evidence for an effect of the treatment on emergence time of bees and bees have not been found to emerge earlier or later from disturbed compared to undisturbed soils (Fig. 3, Appendix S4). There is however a sex difference, well-known as protandry, with males generally emerging earlier than females (Appendix S3). Males emerge on average three days earlier than female bees across all treatments, but there is no evidence that this was altered by soil disturbance.

**Figure 3.**
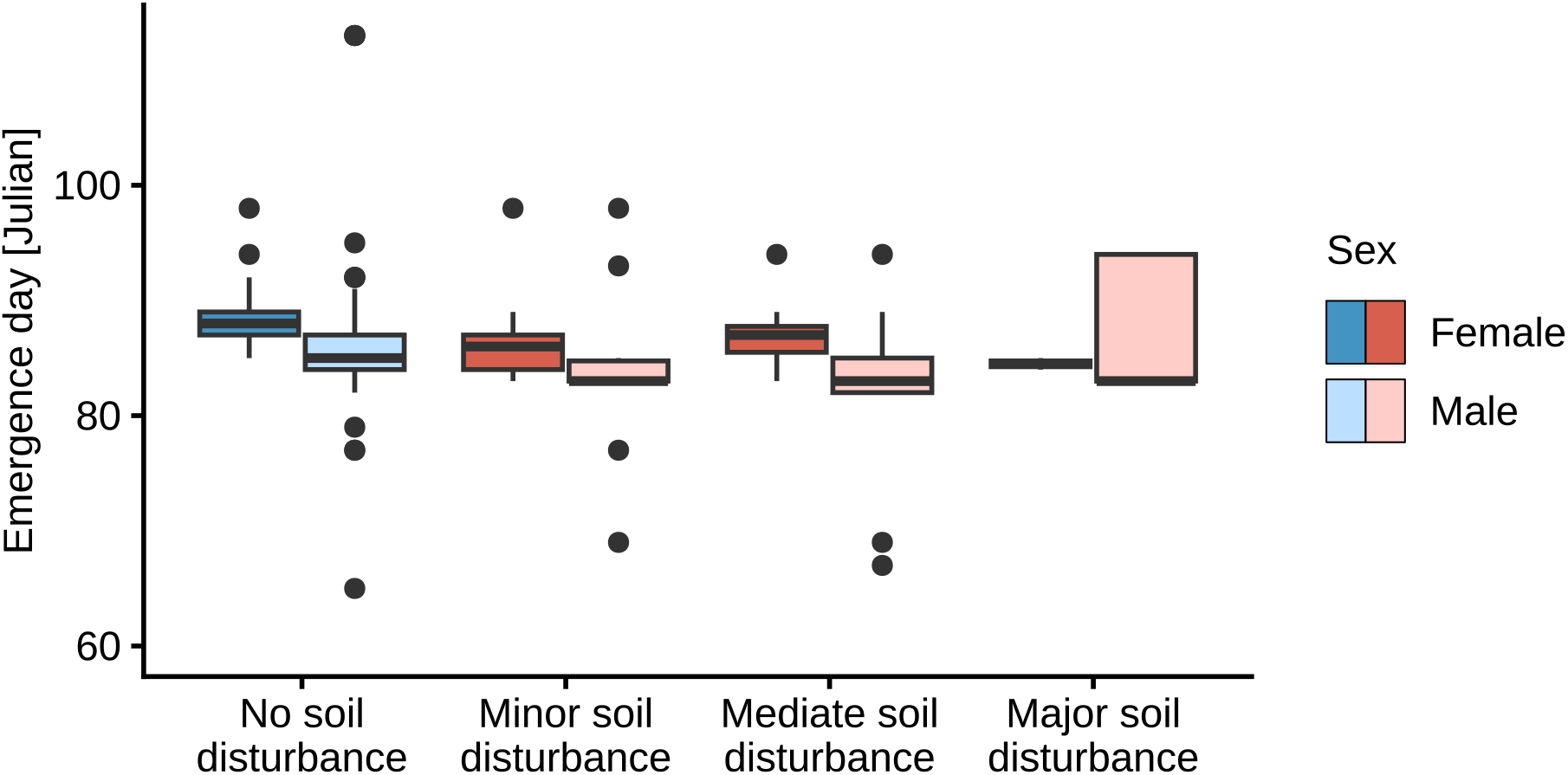
: Bee emergence (on a Julian day scale) distinguishing female and male bees and the four soil disturbance treatments: No, minor, mediate and major soil disturbance.

## Discussion

We investigated the effects of soil translocation and disturbance on the survival and spring emergence time of a solitary ground-nesting bee species. Both translocation and additional disturbance reduced the number of spring mining bees emerging after hibernation. Nevertheless, there were individuals surviving in all treatments, including heavily disturbed soils. Regarding emergence time of bees in spring and protandry (males emerging earlier than females), we found no evidence for an effect of soil translocation and disturbance, and bee emergence time was similar in all soil treatments.

Bees face natural challenges while overwintering, such as frost, desiccation, mould or predation (Keaveny *et al*., 2022; Botsch *et al*., 2024; Cane, 2021). Survival rates vary depending on environmental conditions and on the species. Some species are more resilient than others. In our study, we have used a Colletes species that protects brood cells with a cellophane-like lining strongly protecting against water and bacteria (Almeida, 2008; Antoine and Forrest, 2021; Danforth *et al*., 2019). Individuals were disturbed after full metamorphosis and imagos were translocated in their protecting brood cells. Translocation of larval stages might lead to higher mortalities as larvae are more vulnerable and less well protected against disturbances (Hayes and LÓpez-Martínez, 2021).

In addition to these natural challenges, overwintering survival may be affected by human-induced soil disturbance, including agricultural practices such as ploughing, or soil translocation. Bees and other ground-living insects can survive soil disturbance, but usually with reduced numbers after disturbance (Ullmann *et al*., 2020; Roger-Estrade *et al*., 2010; de Graaff *et al*., 2019). Our results showed that soil disturbance reduced bee survival, with a tendency toward fewer survivors in heavily disturbed soils. Some individuals survived even the severe disturbance and soil translocation treatments. This indicates that although soil disturbance can substantially reduce survival, some individuals may withstand such disturbance, suggesting that translocation of soil containing hibernating bees might be a promising restoration and conservation tool for example in urban areas with construction pressure (Vécrin and Muller, 2003).

Translocation for conservation has been applied predominantly to vertebrates, with comparatively few studies focusing on invertebrates, such as bees and other insects (Taylor *et al*., 2017). The small size and rapid reproduction of insects make them promising candidates for translocation (Thomas *et al*., 2009; Baur *et al*., 2017). Successful insect translocation depends on knowledge of species-specific habitat requirements and reactions to the potential disturbances related to the translocation (Bellis *et al*., 2019). Translocation has a considerable potential to support restoration efforts, particularly for rare species, despite the losses of individuals occurring during the translocation (Osgood, 1974). In our study, translocation reduced survival from an average of 13 to 2 individuals per soil cube, showing that some albeit few individuals survived both translocation and additional soil disturbance treatments. These results suggest that, despite associated survival costs, translocation of soil containing hibernating bees is feasible.

Bee population dynamics and conservation outcomes may also be influenced by behavioural traits such as emergence time. Spring temperature affects emergence time (Fründ *et al*., 2013) and soil disturbance and translocation can expose hibernating bees to altered thermal conditions by changing the depth at which brood cells are located and modifying the surrounding microclimatic conditions. We found no evidence that soil disturbance and translocation affected emergence time. The well-known pattern of protandry remained consistent across treatments, with males emerging earlier than females (Schäffler and Dötterl, 2011). This contrasts with findings from aboveground, cavity-nesting bees, where sex-specific responses to environmental conditions, such as an increase in temperature or season, have been reported (Fründ *et al*., 2013; Wittmann *et al*., 2023).

Soil translocation with varying levels of disturbances might be a promising conservation tool for translocating bee populations during hibernation. High soil disturbances might not be a problem if the nest density of the source population was high, as in our case for the gregariously nesting *C. cunicularius*. Translocating soil is likely more effective than translocating bees during the adult life stage, as bees tend to exhibit strong nest-site fidelity once nesting has started. Our results demonstrate that soil translocation and soil disturbance reduce the survival probability of individuals, but can successfully translocate a proportion of bees while they overwinter within their nesting environment. Such knowledge will become increasingly important as regional to global restoration goals intersect with growing land-use demands from an expanding human population, requiring habitat planning to enhance ecological connectivity.

## Acknowledgements

ACM was supported by the Ministry Of Science, Research and the Arts Baden-Württemberg, Germany from 2021 to 2023. We thank the nature conservation department of the city of Freiburg for necessary research permits and students for their support during field work.

## Data availability

Data and RCode will be made available after publication under doi: https://doi.org/10.5281/zenodo.22735952

## Supporting information

### Supplementary material for

### Ground-nesting bee survival after soil translocation and disturbance

Authors: Anne-Christine Mupepele, Alexandra-Maria Klein, Felix Fornoff

**Table S1:** Linear model summary (M1:Bee survival ∼Treatment*Sex) and likelihood ratio test on bee survival after soil translocation and disturbance.

| Model | Predictor | df | Sum of Squares | F-value | p-value |
| --- | --- | --- | --- | --- | --- |
| M1: ~Treatment*Sex | Treatment | 3 | 314.3413 | 9.4108 | 0.0012 |
| M1: ~Treatment*Sex | Sex | 1 | 1.6652 | 0.1496 | 0.7048 |
| M1: ~Treatment*Sex | Treatment:Sex | 3 | 0.2925 | 0.0088 | 0.9988 |
| M2: ~Treatment+Sex | Treatment | 3 | 314.3413 | 11.4060 | 0.0002 |
| M2: ~Treatment+Sex | Sex | 1 | 1.6652 | 0.1813 | 0.6756 |
| M3: ~Treatment | Treatment | 3 | 314.3413 | 11.9495 | 0.0002 |
| M1:M2 Likelihood Ratio Test | NA | -3 | -0.2925 | 0.0088 | 0.9988 |
| M2:M3 Likelihood Ratio Test | NA | -1 | -1.6652 | 0.1813 | 0.6756 |

**Table S2:**
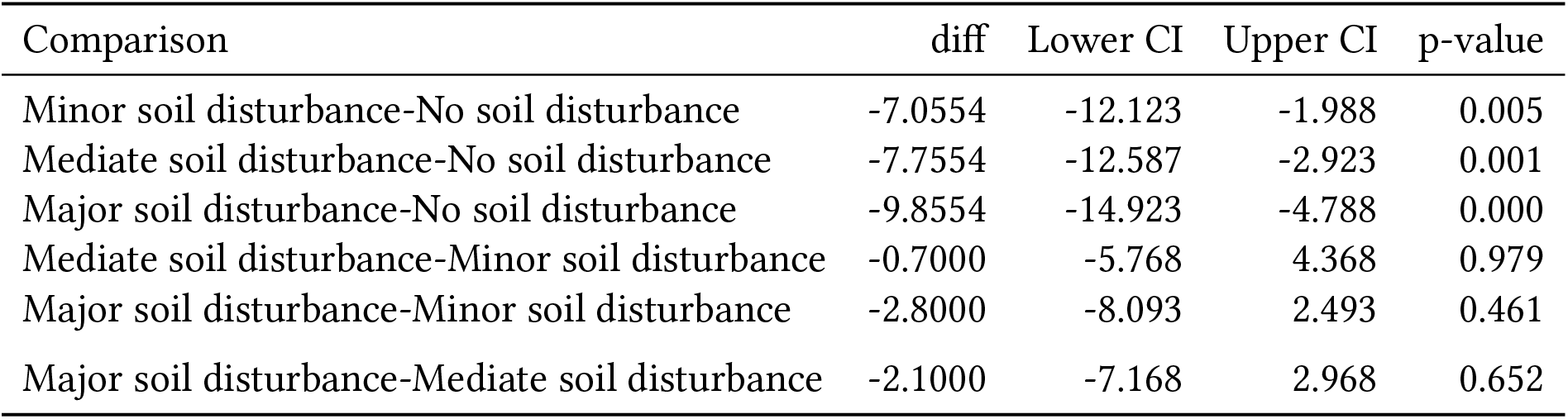
Post-hoc pairwise comparison (TukeyHSD) for the linear model testing how bee survival is influenced by soil disturbance treatments (M3: Bee survival ∼Treatment). Diff= differences in the observed means, Lower CI and Upper CI = the lower and upper end of the confidence interval, p-value= p-value after adjustement for multiple comparisons

| Comparison | diff | Lower CI | Upper CI | p-value |
| --- | --- | --- | --- | --- |
| Minor soil disturbance-No soil disturbance | -7.0554 | -12.123 | -1.988 | 0.005 |
| Mediate soil disturbance-No soil disturbance | -7.7554 | -12.587 | -2.923 | 0.001 |
| Major soil disturbance-No soil disturbance | -9.8554 | -14.923 | -4.788 | 0.000 |
| Mediate soil disturbance-Minor soil disturbance | -0.7000 | -5.768 | 4.368 | 0.979 |
| Major soil disturbance-Minor soil disturbance | -2.8000 | -8.093 | 2.493 | 0.461 |
| Major soil disturbance-Mediate soil disturbance | -2.1000 | -7.168 | 2.968 | 0.652 |

**Table S3:** GLMM summary (Emergence day [Julian day] ∼ Treatment+Sex+randomeffect trap)

| Model | Term | Estimate | Std.Error | z-value | p-value |
| --- | --- | --- | --- | --- | --- |
| M2.1: ~Treatment+Sex | (Intercept) | 4.49 | 0.01 | 694.68 | 0 |
| M2.1: ~Treatment+Sex | Treatment:Minor | -0.02 | 0.01 | -1.79 | 0.07 |
| M2.1: ~Treatment+Sex | Treatment:Mediate | -0.03 | 0.01 | -2.37 | 0.02 |
| M2.1: ~Treatment+Sex | Treatment:Major | 0.00 | 0.02 | -0.08 | 0.94 |
| M2.1: ~Treatment+Sex | Sex:male | -0.03 | 0.01 | -3.94 | 0 |
| M2.1: ~Treatment+Sex | R2_marginal | 0.10 |  |  |  |
| M2.2: ~Sex | (Intercept) | 4.47 | 0.01 | 553.57 | 0 |
| M2.2: ~Sex | Sex:male | -0.03 | 0.01 | -3.88 | 0 |
| M2.2: ~Sex | R2_marginal | 0.06 |  |  |  |

**Table S4:** Likelihood ratio test for model comparison testing the effect of Treatment and Sex on bee emergence time.

| Model | df | Chisq-value | p-value |
| --- | --- | --- | --- |
| M2.1:M2.2 Likelihood Ratio Test | 3 | 5.233037 | 0.155507 |

**Figure S1:**
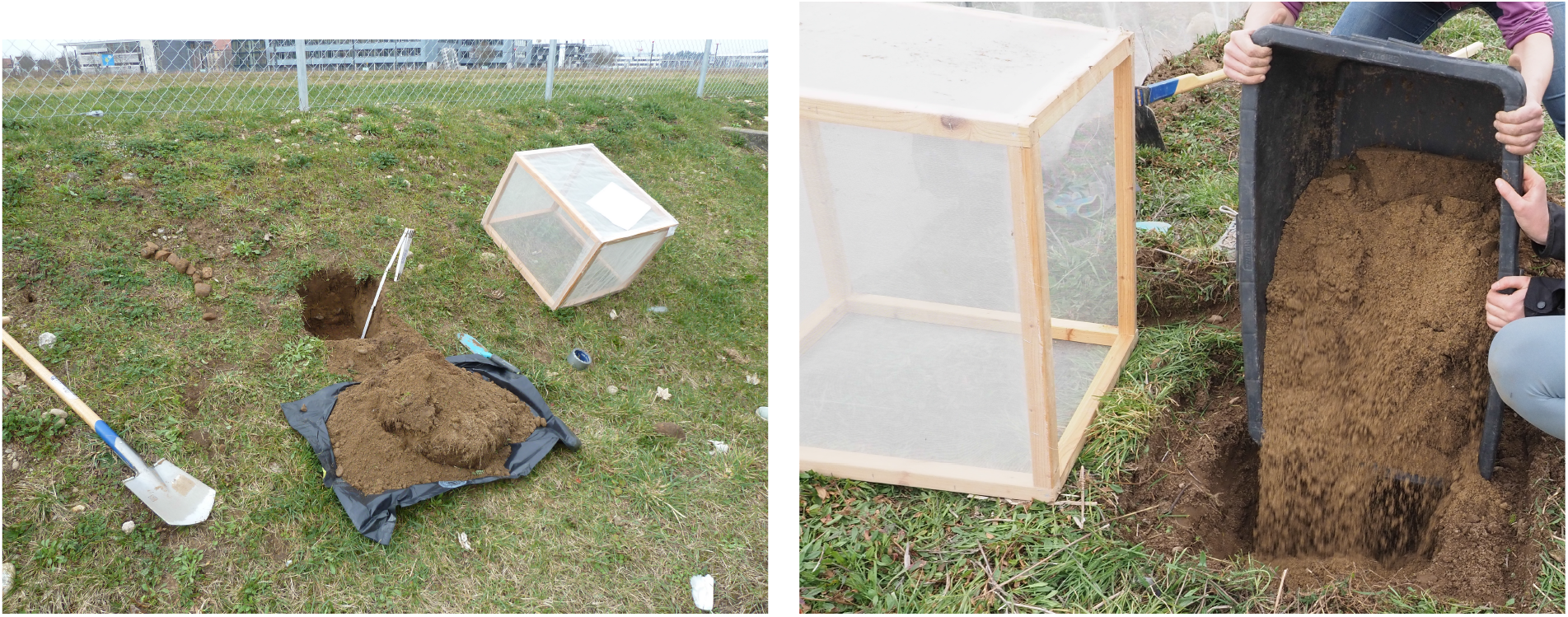
Soil cube excavation at a *C. cunicularius* nest aggregation (left picture). Translocating soil with different disturbance treatments into pre-dug holes (right picture).

S6. RCode and Data All data and R code to reproduce the analysis will be deposited in Zenodo after publication (https://doi.org/10.5281/zenodo.22735952), including an html file of the output, and additionally attached to this manuscript to allow exact reproduction of the analyses.

## Notes

### Competing Interest Statement

The authors have declared no competing interest.

